# Ddx3x regulates B-cell development and immunoglobulin rearrangement in mice

**DOI:** 10.1101/452086

**Authors:** Ke Liu, Durga Krishnamurthy, Jasmine Tuazon, Eshana Mukhopadhyay, Harsha Seelamneni, Erik P. Karmele, Omer A. Donmez, Thomas Perlot, Michelle Foong-Sobis, Rebekah A. Karns, Malay Mandal, Damien Reynaud, Leah C. Kottyan, Matthew T. Weirauch, Wen-Hai Shao, R. Hal Scofield, Josef M. Penninger, John B. Harley, Stephen N. Waggoner

## Abstract

The X chromosome gene, *DDX3X*, is an ATP-dependent RNA helicase with roles in transcription, splicing, nuclear export, and translation. Loss of function mutations in *DDX3X* are linked to a variety of neoplasms, including B-cell lymphoma. We find that conditional homozygous deletion (*Mb1-Cre*) of *Ddx3x* in developing mouse B cells in female mice results in a complete absence of mature peripheral B cells associated with an absolute block at the pro-B cell stage of development in the bone marrow. In male mice with *Vav1-Cre* or *Mb1-Cre* mediated hemizygous deletion of *Ddx3x*, there are less severe reductions in peripheral B-cell frequencies with skewing towards the marginal zone lineage, suggesting that the Y chromosome homolog *Ddx3y* or other male factors may partially compensate for loss of *Ddx3x*. Loss of *Ddx3x* in male mice is associated with perturbations at developmental time points linked to cell cycle arrest and immunoglobulin chain rearrangement. Mechanistically, loss of *Ddx3x* in pre-B cells is associated with reduced expression of the histone reader *Brwd1*, failure to curtail proliferation, and defective *Igk* rearrangement, which skews the peripheral B cell receptor repertoire toward lambda light chain usage. These data reveal that *Ddx3x* plays an essential role in B-cell development by supporting proliferative and epigenetic changes necessary for rearrangement of immunoglobulin genes.

## Introduction

RNA helicases are key regulators of gene expression that control almost all aspects of RNA metabolism from synthesis to degradation (1). Members of the largest group of RNA helicases, the DEAD (Asp-Glu-Ala-Asp) box family, contain not only the characteristic DEAD-box but also a helicase-C domain (2-4). DDX3X (DEAD-Box Helicase 3, X-linked) is a ubiquitously expressed helicase involved in several steps of RNA processing that ultimately promote translation (5-9). These functions contribute to roles for DDX3X in cell cycle control, apoptosis, innate immunity, and tumorigenesis (10-14). (10-14). The amino acid sequence homology between human and mouse DDX3X is >98%, revealing strong evolutionary conservation that likely reflects conserved function (15). Of note, *DDX3X* has a homolog on the Y chromosome, *DDX3Y* (91% identity), which exhibits overlapping helicase activity but that is putatively only expressed in testes (16-19). However, isoform specific antibodies reveal that DDX3Y is expressed in male mouse B cells and that DDX3Y expression is increased in *Ddx3x*-deficient tumors (20).

DDX3X is broadly expressed in hematopoietic cells (21) and mutations in *DDX3X* are linked to various cancers (3, 9), including leukemia and lymphoma. Yet, the functions of DDX3X in normal development of immune cells remain incompletely defined. Global deletion of *Ddx3x* is embryonically lethal (22), and deletion of *Ddx3x* in hematopoietic cells perturbs erythropoiesis to cause early developmental arrest (20). We and others find that hemizygous loss of *Ddx3x* in hematopoietic cells negatively impacts the development of B cells (20, 23, 24).

By generating mice where *Ddx3x* is conditionally and efficiently deleted at early stages of B-cell differentiation using *Mb1-Cre*, we demonstrate that *Ddx3x* is essential for generation of a B cell compartment in female mice. This finding expands on recent observations with *Cd19-Cre*-mediated deletion of *Ddx3x* at later stages of B-cell development (20). In contrast to our recent observation that *Ddx3y* could not compensate for loss of *Ddx3x* in the development of mature NK cells (25), we found that a diminished but functional B-cell compartment could be generated in hemizygous *Ddx3x*-deficient (*Mb1-Cre*) male mice. Evaluations of surviving B cells in the bone marrow and spleens of these hemizygous mice reveal alterations in the peripheral repertoire of B cells as well as dysregulation of proliferative arrest and immunoglobulin gene rearrangement at key developmental checkpoints.

## RESULTS

### Ddx3x is essential for B-cell development in female mice

Previous work showed that *Cd19-Cre*-mediated deletion of both copies of *Ddx3x* during B cell development in the bone marrow of female mice results in reduced numbers of immature and mature B cells (20). We hypothesized that the low recombination efficiency of *Cd19-Cre* at early stages of B cell differentiation could contribute to the incomplete blockade in B-cell development in that prior study. To efficiently delete *Ddx3x* at earlier development time points when precursors commit to the B-cell lineage, we bred *Ddx3x*^*FL*/*FL*^ mice (23) to *Mb1-Cre* mice (26). The resulting female *Ddx3x*^*ΔMb1*^ animals exhibited a near total absence of B cells in the spleen (**Figure 1A,B**) and bone marrow (**Figure 1C**) relative to littermate controls. This resulted from a complete block in B-cell differentiation at the pro B cell stage of development (**Figure 1D**, fraction B, CD19^+^ B220^+^ CD43^+^ CD24^+^ BP-1^neg^) of development that prevented generation of subsequent fractions of pre B cells, immature B cells, and mature B cells (**Figure 1D-F**). Therefore, *Ddx3x* is required in developing B cells for the generation of a B cell compartment in female mice.

**Figure 1.**
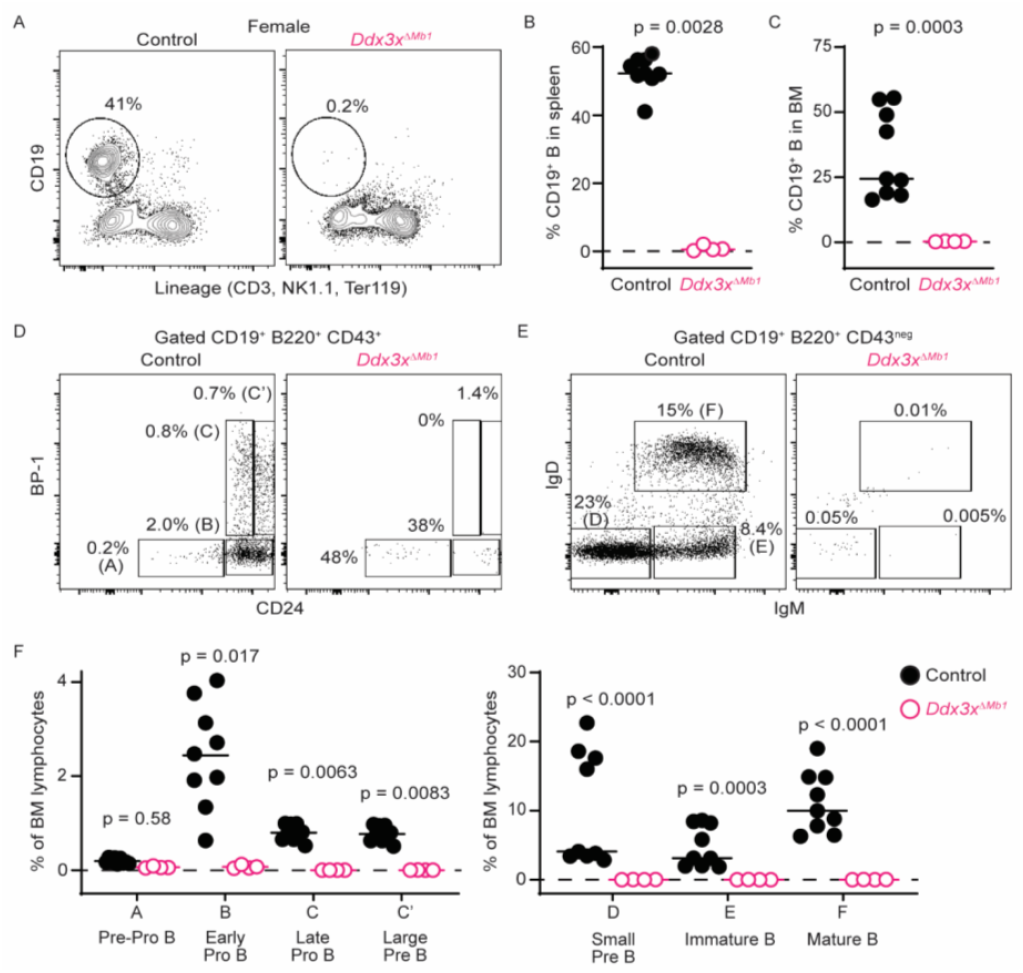
*Ddx3x* is required for B cell development in female mice. (**A,B**) Flow cytometry analysis of splenocytes from female *Ddx3x*^*ΔMb1*^ (n=4) and *Cre*-negative littermate control (n=9) mice showing (**A**) representative contour plots and (**B**) proportions of CD19^+^ lineage^neg^ B cells. (**C**) Corresponding proportions of B cells in bone marrow. (**D-F**) Developmental stages of B cell development in bone marrow, gated based on (**D**) CD24 and BP-1 expression on CD43^+^ B cells or (**E**) surface IgM and IgD on CD43^neg^ B cells. Horizontal dotted lines are placed at y=0. Data are pooled from two independent experiments. Statistical significance determined using (**B**) Mann Whitney U test (non-normal distribution), (**C**) unpaired t test with Welch’s correction (unequal variance), or (**F**) Kruskal-Wallis test (non-normal distribution) with multiple comparisons controlled for false discovery rate using a two-stage linear step-up procedure.

### Ddx3y mitigates loss of B cells in male mice lacking Ddx3x

We recently found that ablation of *Ddx3x* during NK cell development results in deficiency of NK cells in both male and female mice (25), suggesting that *Ddx3y* is unable to compensate for *Ddx3x* in that lymphocyte lineage. In contrast, the presence of *Ddx3y* mitigated the loss of B cells in male *Ddx3x*^*ΔCd19*^ relative to their *Ddx3x*-deficient female counterparts (20). Consistent with that observation, we observed a ∼3-fold reduction in peripheral B cell counts but maintenance of more than 20 million B cells in male *Ddx3x*^*ΔMb1*^ mice (**Figure 2A,B**). *Cre*-mediated deletion of the floxed *Ddx3x* allele was robust in surviving splenic B cells (**Figure 2C**). Reduced frequencies of B cells were also observed in the spleen and inguinal lymph node of male mice in which *Ddx3x* was deleted in all hematopoietic cells via *Vav1-Cre* (*Ddx3x*^*ΔVav1*^, **Figure 2D**). Thus, the stringent requirement for *Ddx3x* in B cell development in female mice is not shared by loss of the single copy of *Ddx3x* in male mice, possibly due to the presence of *Ddx3y*, which is known to be expressed in mature B cells, is over-expressed upon loss of *Ddx3x*, and partially rescues lymphogenesis (20). Other Y chromosome-restricted genes, or other ill-defined male-specific factors also may contribute.

**Figure 2.**
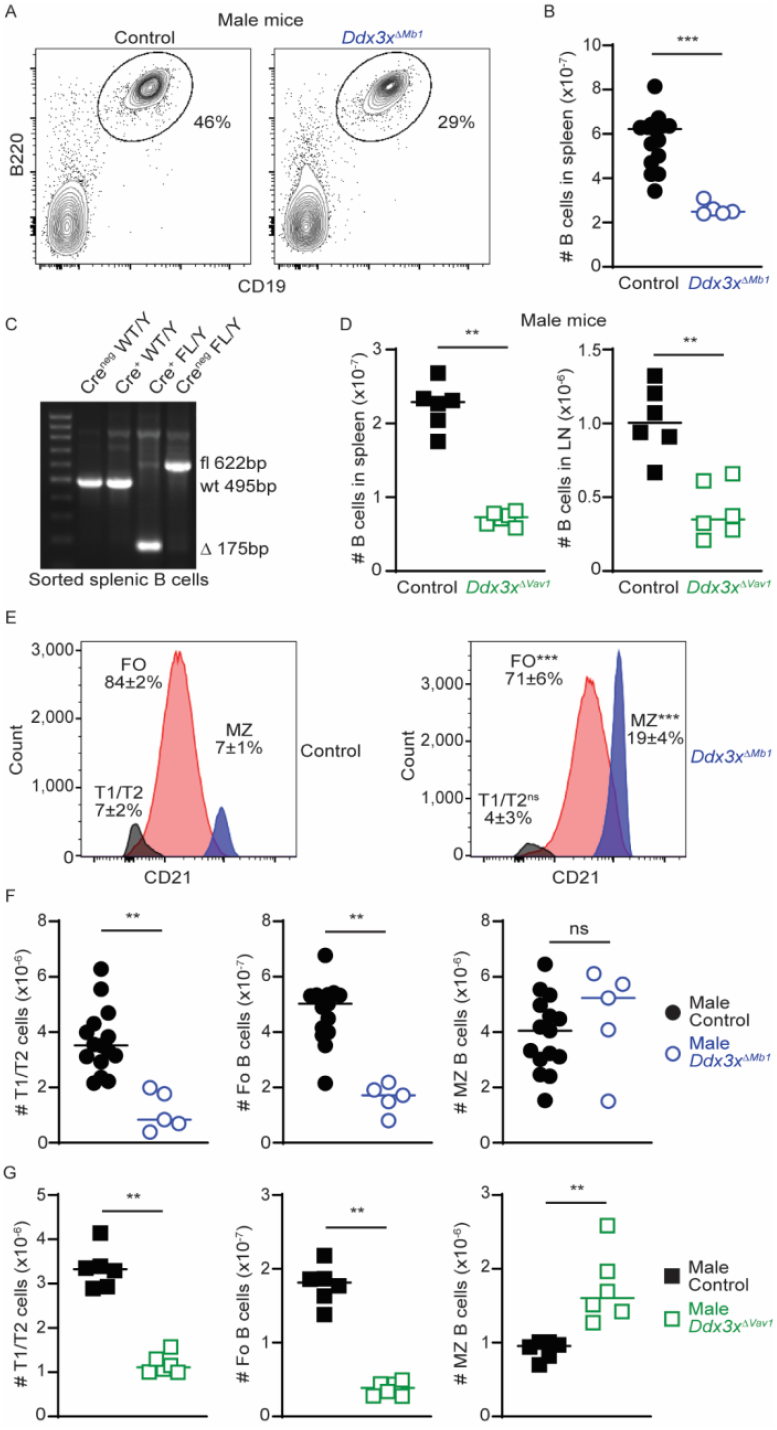
Reduced B cell development with hemizygous loss of *Ddx3x* in male mice. (**A,B**) Flow cytometry analysis of splenocytes from male *Ddx3x*^*ΔMb1*^ (n=5) and *Cre*-negative littermate control (n=15) mice showing (**A**) representative contour plots and (**B**) proportions of CD19^+^ lineage^neg^ B cells. (**C**) *Ddx3x* genotyping PCR with sorted splenic B cells from wild-type male *Mb1-Cre*+ or *Cre*^neg^ mice (495bp wild-type product), control *Cre*^neg^ *Ddx3x*^*FL/Y*^ (622bp floxed product), and *Cre*^+^ *Ddx3x*^*FL/Y*^ (175bp deletion (Δ) product). (**D**) Proportions of B cells in spleen (left) and inguinal lymph node (LN, right) of male *Ddx3x*^*ΔVav1*^ (n=6) and *Cre*-negative littermate controls (n=6). (**E**) Representative overlay with mean (±SD) proportions and corresponding (**F**) numbers of transitional (T1/T2, IgM^high^ CD21^neg^), follicular (FO, IgM^low^ CD21^+^ CD23^+^), and marginal zone (MZ, CD21^hi^ CD1d^hi^ CD23^neg^) in male *Ddx3x*^*ΔMb1*^ (n=5) and *Cre*-negative littermate control (n=15) mice. (**G**) Numbers of each splenic B cell subset in male *Ddx3x*^*ΔVav1*^ (n=6) and *Cre*- negative littermate control (n=6) mice. Data are pooled from 2 -3 independent experiments. Statistical significance was determined using a Mann Whitney U test (ns, p>0.1; **, p<0.01; ***, p<0.001).step-up procedure.

### Loss of Ddx3x skews B cell differentiation to marginal zone phenotypes

Examination of the B cells that are present in the spleen of male *Ddx3x*^*ΔMb1*^ mice revealed a marked skewing towards a CD21^high^ CD1d^high^ marginal zone (MZ) phenotype (**Figure 2E**). Consequently, the diminished number of B cells in male *Ddx3x*^*ΔMb1*^ mouse spleens is a consequence of reductions in the transitional (T1/T2) and follicular (FO) B cell compartments (**Figure 2F**). We observed a similar skewing towards MZ phenotypes among B cells in male *Ddx3x*^*ΔVav1*^ mice (**Figure 2G**). Considered alongside observations in *Ddx3x*^*ΔCd19*^ mice (20), these observations reinforce that peripheral B cell differentiation is skewed by loss of *Ddx3x* in a B-cell intrinsic manner.

### B-cell extrinsic factors drive hypergammaglobulinemia following hematopoietic loss of Ddx3x

The skewed differentiation of peripheral B cells in male mice lacking *Ddx3x* led us to assess changes in circulating antibodies. Indeed, hemizygous deletion of *Ddx3x* in hematopoietic cells (*Ddx3x*^*ΔVav1*^) resulted in a 2-to 4-fold increase in sera IgM and IgG relative to littermate controls (**Figure 3A**). In contrast, sera IgA titers were relatively unchanged and IgE levels were decreased nearly 9-fold in male *Ddx3x*^*ΔVav1*^ **(Figure 3A)**. Among IgG subclasses, IgG1, IgG2b, and IgG3 titers were increased in male *Ddx3x*^*ΔVav1*^ mice while IgG2c titers remained unaltered (**Figure 3B**). However, we did not observe any changes in autoantibody titers over the first year of life in in male *Ddx3x*^*ΔVav1*^ mice compared to littermate controls (**Figure 3C**). These data reveal hypergammaglobulinemia in *Ddx3x*^*ΔVav1*^ mice despite clear deficiencies in the quantity of B cells.

**Figure 3.**
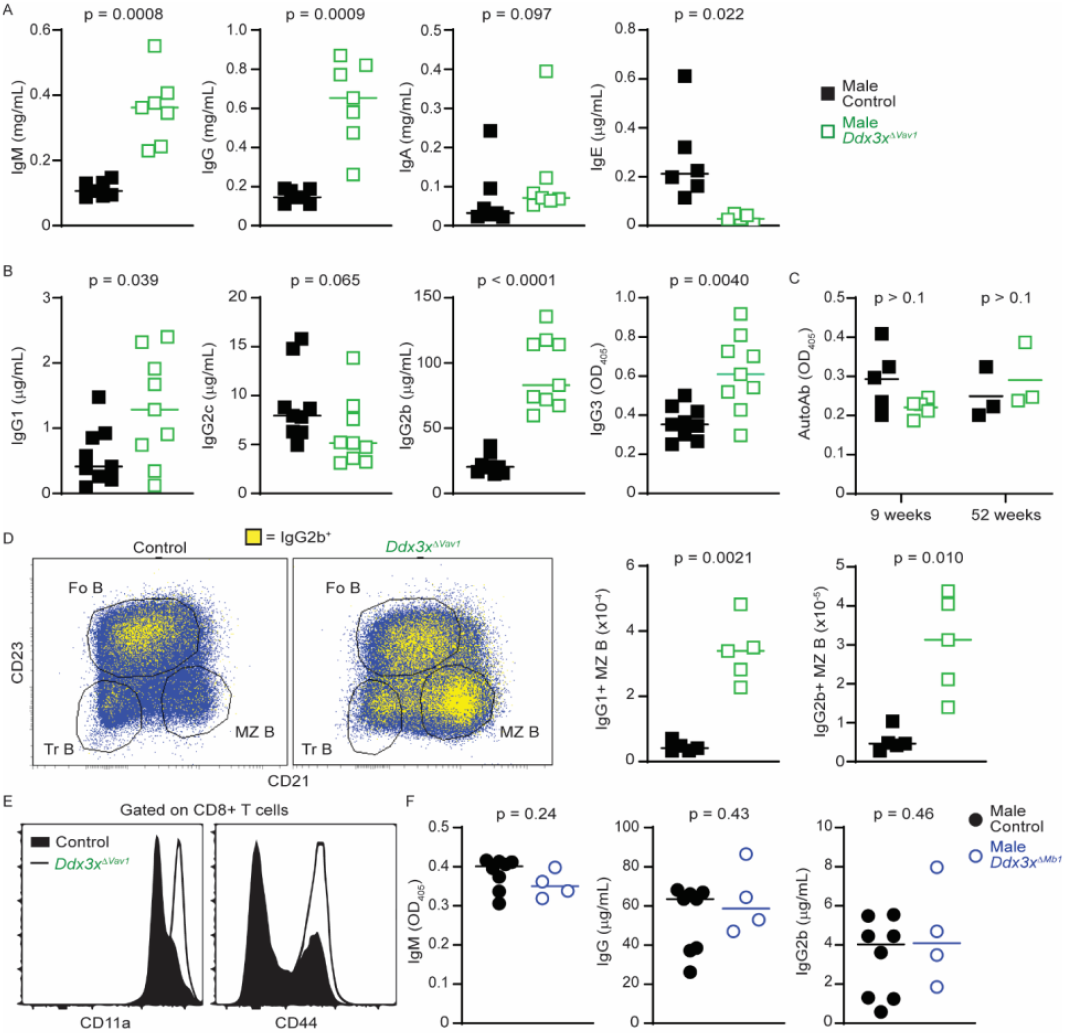
B-cell extrinsic hypogammaglobulinemia caused by *Vav1*-driven *Ddx3x* deletion. ELISA measurement of (**A-B**) immunoglobulin titers in sera of male *Ddx3x*^*ΔVav1*^ (n=7) and *Cre*- negative littermate control (n=6) mice. (**C**) Autoantibody titers in sera of male *Ddx3x*^*ΔVav1*^ (n=3-4) and control (n=3-6) mice at 9 or 52 weeks of age. (**D**) Representative flow cytometry overlays (left) of IgG2b+ (yellow) B cells among total (blue) separated into transitional (Tr), follicular (Fo), and marginal zone (MZ) phenotypes based on CD21 and CD23 expression. Right panels show quantification of increased numbers of IgG1+ or IgG2b+ MZ B cells in male *Ddx3x*^*ΔVav1*^ and control mice (n=5/group). (**E**) Histogram overlays showing representative altered CD11a or CD44 expression phenotype of gated CD8 T cells in male *Ddx3x*^*ΔVav1*^ and control mice. (**F**) ELISA measurement of immunoglobulin titers in sera of male *Ddx3x*^*ΔMb1*^ (n=4) and *Cre*-negative littermate control (n=8) mice. Data are pooled from two independent experiments. Significant differences were calculated using a (**A,B,D,F**) Mann Whitney U test or a two-tailed unpaired t test with Welch’s correction, depending on the normality of the data distribution. A Kruskal-Wallis test with multiple comparisons controlled for false discovery rate using two-stage linear step-up procedure was applied to part (**C**).

We hypothesized that the skewed marginal zone (MZ) phenotype of *Ddx3x*-deficient B cells may contribute to exaggerated production of immunoglobulins. Indeed, while few marginal zone B cells scored positive for IgG2b or IgG1 in control mice, there was a marked increase in the proportion and number of MZ B cells positive for either IgG2b (shown in representative dot plots) or IgG1 in *Ddx3x*^*ΔVav1*^ mice (**Figure 3D**). These responses were associated with a marked lymphopenia in *Ddx3x*^*ΔVav1*^ mice (data not shown) that likely contributes to homeostatic proliferation and inflammation. Indeed, CD8 T cells (**Figure 3E**) and CD4 T cells (data not shown) in the spleens of *Ddx3x*^*ΔVav1*^ mice exhibit phenotypes consistent with lymphopenia-induced homeostatic proliferation (28), including elevated expression levels of activation markers CD44 and CD11a (**Figure 3E**).

To test whether inflammatory lymphopenia caused by hematopoietic loss of *Ddx3x* is a B-cell extrinsic driver of these phenotypes, we measured immunoglobulin titers in mice with B-cell restricted deletion of *Ddx3x* (*Ddx3x*^*ΔMb1*^ mice). In contrast to *Ddx3x*^*ΔVav1*^ mice (**Figure 3A,B**), there were no elevations in IgM, IgG, or IgG2b in male *Ddx3x*^*ΔMb1*^ mice (**Figure 3F**). Thus, skewed MZ differentiation observed in each of these strains (**Figure 2E-G**) does not directly cause hypergammaglobulinemia, which instead is an apparent consequence of lymphopenia-induced inflammation unique to *Ddx3x*^*ΔVav1*^ mice.

### Ddx3x supports B-cell lymphopoiesis

The peripheral reductions in B cell numbers and skewed marginal zone differentiation in *Ddx3x*^*ΔVav1*^ and *Ddx3x*^*ΔMb1*^ mice portends a defect in B-cell development in the bone marrow. In contrast to the absence of B cells in the bone marrow of female *Ddx3x*^*ΔMb1*^ mice (**Figure 1C**), male *Ddx3x*^*ΔMb1*^ mice exhibit relatively normal frequencies of B cells in the bone marrow (**Figure 4A**). We examined stage-specific progression of B-cell development in the bone marrow of male *Ddx3x*^*ΔMb1*^ and control mice (**Figure 4B**). In contrast to the early pro B cell block seen in female *Ddx3x*^*ΔMb1*^ mice (**Figure 1F**), there were statistically similar proportions (**Figure 4C**) and numbers (**Figure 4D**) of pre-pro, pro, and large pre B cells in male *Ddx3x*^*ΔMb1*^ and control mice. A small reduction in early pro B cells frequencies was apparent in male *Ddx3x*^*ΔMb1*^ mice that did not reach statistical significance (**Figure 4C**). Yet, a significant reduction in the proportions of small pre B cells and immature B cell was apparent in *Ddx3x*^*ΔMb1*^ mice (**Figure 4C**) that was not mirrored by a significant reduction in cell counts (**Figure 4D**).

**Figure 4.**
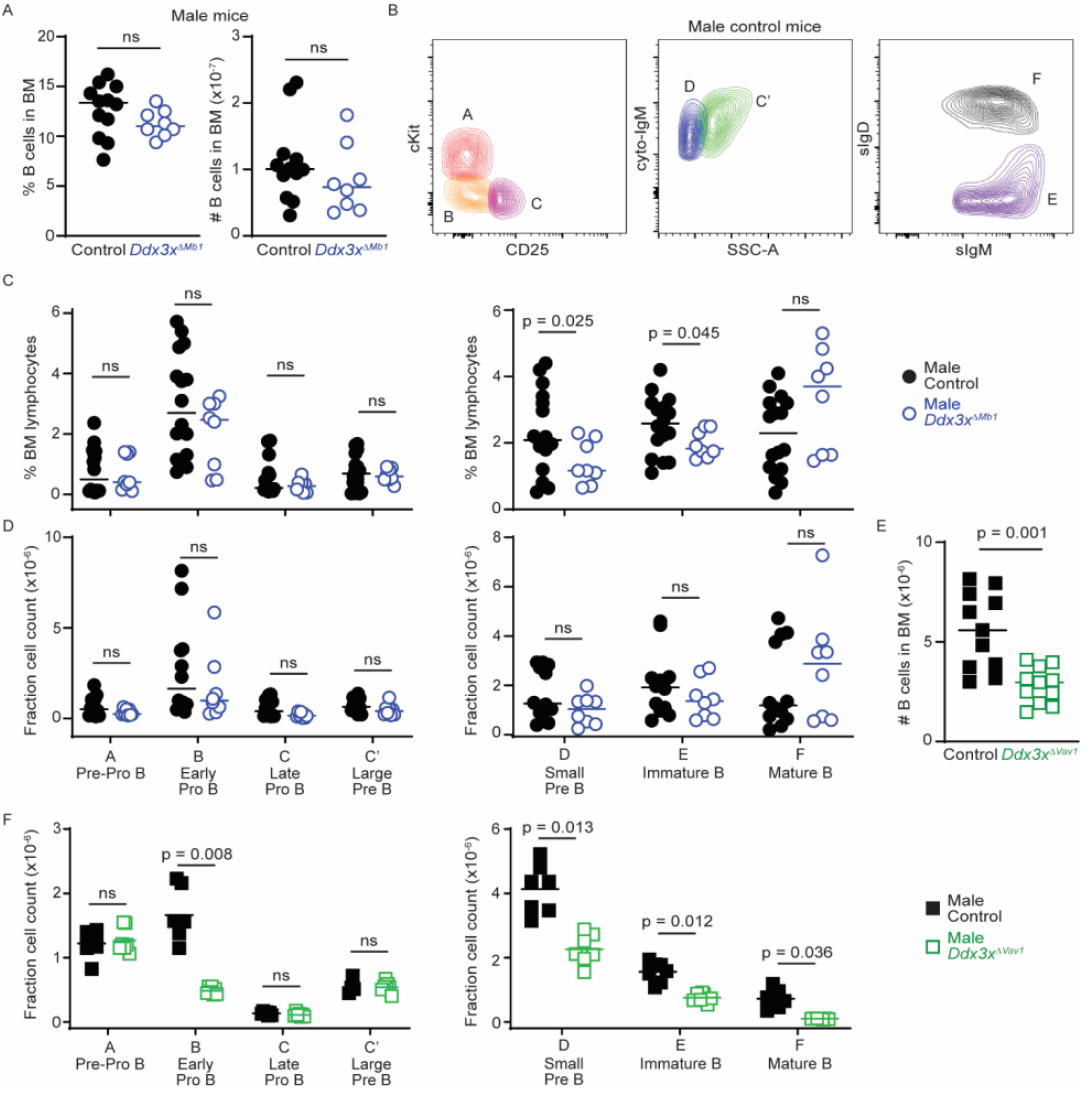
Hemizygous loss of *Ddx3x* causes partial blocks at early pro-B and small pre-B stages of B cell development. (**A**) Flow cytometry quantification of CD19^+^ lineage^neg^ bone marrow B cells in male *Ddx3x*^*ΔMb1*^ (n=8) and *Cre*-negative littermate control (n=10) mice. (**B-D**) Developmental stage subsetting of bone marrow B cells as fractions A (pre-pro B, cKit^+^ CD25^neg^ cytoplasmic (cytoIgM)^neg^ surface IgM (sIgM)^neg^ sIgD^neg^), B (early pro B, cKit^neg^ CD25^neg^ cytoIgM^neg^ sIgM^neg^ sIgD^neg^), C (late pro B, cKit^neg^ CD25^+^ cytoIgM^neg^ sIgM^neg^ sIgD^neg^), C’ (large pre B, side-scatter area (SSC-A)^high^ cytoIgM^+^ sIgM^neg^ sIgD^neg^), D (small pre B, SSC-A^low^ cytoIgM^+^ sIgM^neg^ sIgD^neg^), E (immature B, sIgM^+^ sIgD^neg^), and F (mature B, sIgM^+^ sIgD^+^). Representative overlay of gated fractions shown in **B**, with quantification of (**C**) proportions and (**D**) numbers of each subset in male *Ddx3x*^*ΔMb1*^ (n=8) and control (n=10) mice. (**E,F**) Parallel flow cytometry analysis of numbers of B cells and B-cell developmental stages in male *Ddx3x*^*ΔVav1*^ and *Cre*-negative littermate control (n=6/group) mice. Data are pooled from 2-3 independent experiments. Significant differences were calculated using (**A,E**) a two-tailed unpaired t test with Welch’s correction, (**C,D**), **a** Kruskal-Wallis test with multiple comparisons controlled for false discovery rate using a two-stage linear step-up procedure, or (**F**) a Brown-Forsythe ANOVA test with Dunnett’s T3 multiple comparisons test. ns, p>0.1.

A similar but exaggerated pattern was observed in male *Ddx3x*^*ΔVav1*^ mice, which exhibited fewer bone marrow B cells (**Figure 4E**) corresponding to signficant throttling of B cell development at the early pro B cell and small pre B cell stages. Of note, the frequencies of late and large pro-B cells were normal (**Figure 4E**), with similar expression levels of intracellular IgM (*Ddx3x*^*ΔVav1*^ *=* 47.2 ± 1.6%, WT = 47.5 ± 1.3%, p=0.90, n=4, two-tailed unpaired t-test), suggesting B cells progressing through early pro-B cell stage properly rearranged heavy chain. In male *Ddx3x*^*ΔVav1*^ mice, the defect arises at the small pre B cell stage, persisting through the immature and mature B cells stages (**Figure 4E**), likely contributing to peripheral B-cell lymphopenia. Overall, these results suggest that DDX3X is important at B cell developmental stages linked to cell cycle arrest and immunoglobulin chain rearrangement. Male mice partially compensate for loss of *Ddx3x* to a degree that is dependent on the developmental timing of *Ddx3x* deletion in hematopoietic progenitors (*Vav1-Cre*), early B cell development (*Mb1-Cre*), and later B cell development (*Cd19-Cre*, (20)).

### Ddx3x supports Brwd1 expression and Igκ rearrangement in male pre-B cells

Perturbations of B cell development associated with *Ddx3x* deletion are linked to developmental time points where B cells undergo cell cycle arrest, epigenetic changes, and rearrangement of heavy (early pro B) and light (small pre B) immunoglobulin chains. The extreme paucity of B cells in female *Ddx3x*^*ΔMb1*^ mouse bone marrow complicated exploration of underlying mechanisms of Ddx3x regulation of B cell development. Therefore, we focused on the major perturbation at the pre B cell stage in *Ddx3x*^*ΔVav1*^ male mice that is mirrored to a lesser extent in *Ddx3x*^*ΔMb1*^ male mice.

We extracted RNA from small pre-B cells (CD3^-^Ly6C^-^Ter119^-^CD11b^-^Gr-1^-^B220^+^CD43^-^IgM^-^IgD^-^) sorted out of the bone marrow of male *Ddx3x*^*ΔVav1*^ and WT mice. We then analyzed gene expression using Affymetrix mouse gene 2.0 arrays. We found 477 differentially expressed transcripts that have more than 1.5-fold expression differences between *Ddx3x*^*ΔVav1*^ and control pre-B cells and that reached a threshold (p<0.05) of statistical significance (**Figure 5A**). Transcripts expressed at lower levels in the *Ddx3x*^*ΔVav1*^ small pre-B cells compared to controls included several light chain genes (*Iglc2, Igkv10-96, Iglv1*) and the histone reader *Brwd1. Brwd1* (bromodomain and WD repeat domain containing 1) encodes a protein that targets and restricts recombination at the immunoglobulin *Kappa* (*Igκ)* locus in B cells (27). Notably, *Brwd1* mutant mice exhibited a similar B-cell phenotype to that observed in *Ddx3x*^*ΔVav1*^ mice, with both mouse strains exhibiting a deficiency at the small pre-B cell stage of B-cell development in bone marrow and peripheral B-cell lymphopenia (27). We hypothesized that *Ddx3x* promotes upregulation of *Brwd1* expression at the pre-B cell stage, such that reduced expression of *Ddx3x* causes a decrease in the expression of *Brwd1* and defects in B-cell light chain recombination.

**Figure 5.**
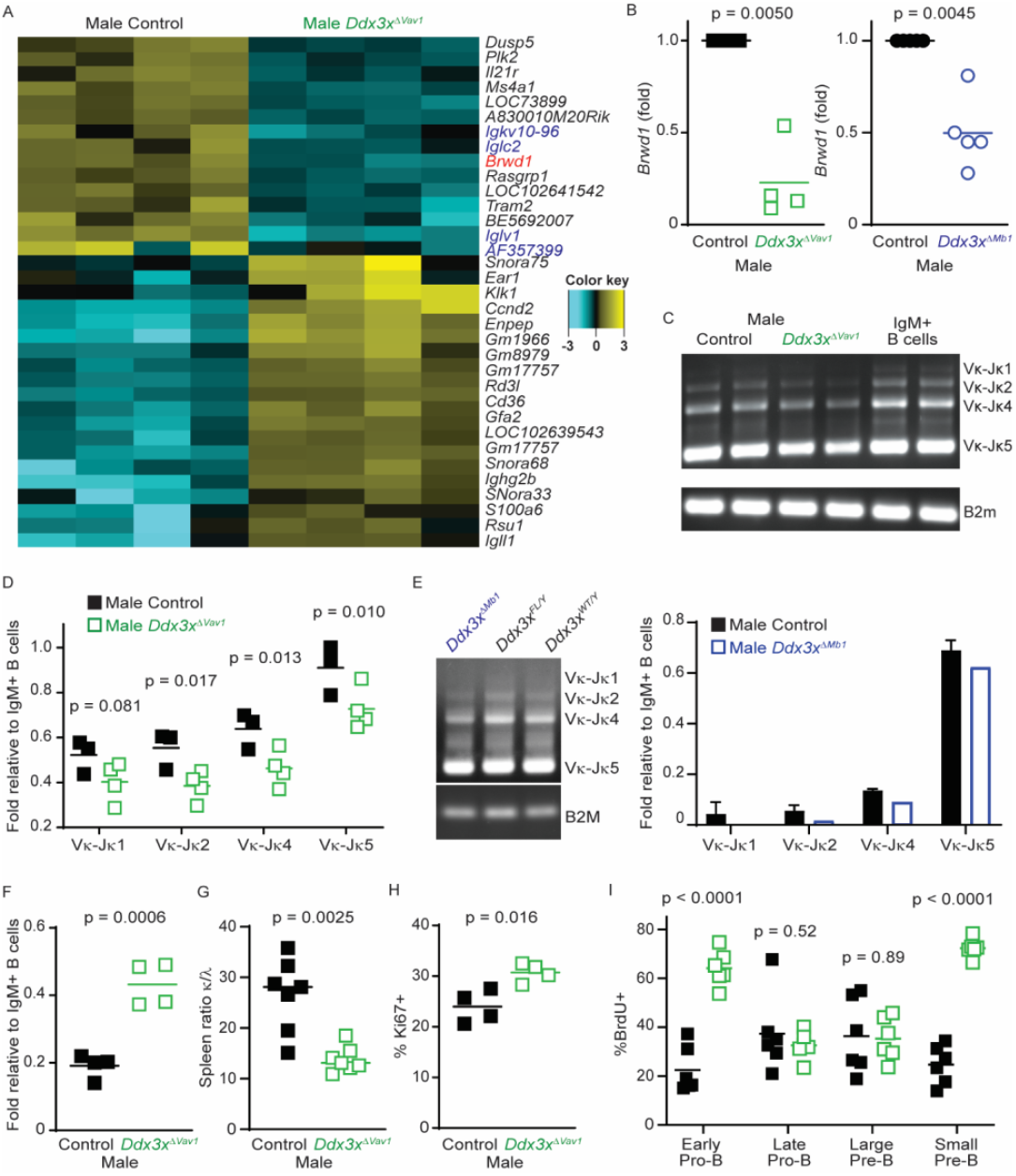
Hemizygous loss of *Ddx3x* perturbs *Brwd1* expression, *Kappa* chain rearrangement, and cell cycle arrest in small pre-B cells. (**A**) Clustered heat map of top 15 up-regulated and down-regulated genes in small pre-B cell from male *Ddx3x*^*ΔVav1*^ mice compared to littermate controls (n=4/group). (**B**) *Brwd1* mRNA expression (qPCR) in small pre-B cell from male *Ddx3x*^*ΔVav1*^ or *Ddx3x*^*ΔMb1*^ mice normalized to levels in littermate controls (n=4-5/group). (**C-E**) Semi-quantitative PCR analysis of *Igk* rearrangement in bone marrow small pre-B cell from (**C,D**) male *Ddx3x*^*ΔVav1*^ and control (n=4/group) or (**E**) *Ddx3x*^*ΔMb1*^ and control (n=1-2/group) mice. Wild-type splenic CD19^+^IgM^+^ cells were used as a positive control. (**D,E**) Quantification of Vκ-Jκ1,2,4,5 band intensity. (**F**) Quantitative real-time PCR analysis of *Vκ-Jκ1* expression in male *Ddx3x*^*ΔVav1*^ and littermate control small pre-B cell normalized to wild-type splenic CD19^+^IgM^+^ cells. (**G**) Ratio of κ to λ light chain usage by male *Ddx3x*^*ΔVav1*^ and control splenic B cells (n=7/group) determined by flow cytometry. (**H**) Bone marrow small pre-B cells (CD19^+^sIgM^-^CD43^-^) were assessed for Ki67 staining in male *Ddx3x*^*ΔVav1*^ and control (n=4/group). (**I**) *In vivo* BrdU incorporation by bone marrow B cell subsets assessed by flow cytometry (n=6/group). Data are pooled from 2-3 independent experiments. Significant differences were calculated using (**B**) a one sample t test; (**D**) an ordinary one-way ANOVA or (**I**) a Brown-Forsythe ANOVA with multiple comparisons controlled for false discovery rate using a two-stage linear step-up procedure; and (**F,G,H**) a two-tailed unpaired t test with Welch’s correction.

We confirmed that the expression of *Brwd1* is decreased in small pre B cells of male *Ddx3x*^*ΔVav1*^ and *Ddx3x*^*ΔMb1*^ mice relative to controls (**Figure 5B**). We also observed that *Ddx3x*^*ΔVav1*^ pre B cells exhibit reduced expression of a germline *Igk* transcript relative to wild-type controls (*Ddx3x*^*ΔVav1*^ *Igk* fold relative to control = 0.63 ± 0.01, p = 0.013). Germline *Igk* transcripts are conventionally upregulated at this stage of development as a consequence of altered chromatin accessibility in the *Kappa* locus prior to DNA recombination (29). We used semi-quantitative PCR to demonstrate that rearrangement of Vκ-Jκ was diminished 20-30% in *Ddx3x*^*ΔVav1*^ small pre B cells compared to control small pre B cells (**Figure 5C, D**). A similar trend was observed in *Ddx3x*^*ΔMb1*^ small pre B cells (**Figure 5E**). Spleen-derived wild-type IgM^+^ B cells served as a positive control for *Kappa* rearrangement in this assay and for qPCR-based confirmation of reduced efficiency of Vκ-Jκ1 rearrangement in *Ddx3x*^*ΔVav1*^ small pre B cells (**Figure 5F**). A consequence of inefficient *κ* light chain rearrangement in *Ddx3x*^*ΔVav1*^ mice manifested as a significant skewing of the ratio of *κ* to *λ* light chain usage in splenic B cells in these mice (**Figure 5G**). These data demonstrate that *κ* light chain rearrangement is deficient in developing male B cells in the absence of *Ddx3x*.

In order to facilitate light chain rearrangement, developing B cells must efficiently exit the cell cycle at the small pre-B cell stage. In conjunction with reduced light chain rearrangement, more pre-B cells in *Ddx3x*^*ΔVav1*^ mice stain positive for Ki67, a marker of active cell cycling (**Figure 5H**). Bromodeoxyuridine (BrdU) incorporation as a marker of progression through S phase was lower in early pro B and small pre B cells compared to late pro B and large pre B cells as a consequence of cell cycle arrest at the former stages (**Figure 5I**). Of note, early pro B and small pre B cells in *Ddx3x*^*ΔVav1*^ mice exhibited a marked increase in BrdU incorporation relative to controls (**Figure 5I**). In summary, these data show a defect in *Brwd1* expression, cell cycle arrest, and *κ* light chain rearrangement in small pre B cells in the absence of *Ddx3x*.

## DISCUSSION

We define a requirement for *Ddx3x* at critical stages of immunoglobulin gene rearrangement to facilitate generation of a peripheral B cell compartment in mice. Our data are consistent with *Ddx3y* partially compensating for loss of *Ddx3x* to support B cell development in male mice. B cell-intrinsic promotion of *Brwd1* expression, cell cycle arrest, and κ light chain recombination defines the mechanistic contributions of *Ddx3x* at the small pre B cell stage of B cell development. Hemizygous loss of *Ddx3x* skewed B cell phenotypes to favor marginal zone differentiation, which contributed to hypergammaglobulinemia driven by B-cell extrinsic inflammation associated with lymphopenia. These results represent a new cellular function for *Ddx3x* in lymphocyte biology.

We contributed key early evidence for B cell defects and embryonic lethality with hematopoietic loss of *Ddx3x* (23, 24). These phenotypes were extended by the Möröy Lab, who were the first to show a B-cell intrinsic role for *Ddx3x* using *Cd19-Cre* (20). The lower efficiency of *Cd19-Cre* at early stages of B cell development predominately manifested as a marked loss of mature B cells in the bone marrow and periphery of female *Ddx3x*^*ΔCd19*^ mice (20). The new data in this manuscript using *Mb1-Cre* to delete *Ddx3x* earlier in B lymphopoiesis provide a critical new insight that *Ddx3x* is absolutely required for progression through the early pro B cell stage of development in female mice.

In addition, our new data reinforce the potential of *Ddx3y* to partially compensate for loss of *Ddx3x* in the development of B cells in male mice. This contrasts sharply with our recent report of a DDX3X-specific necessity for DDX3 helicase activity in NK cells (25). A likely explanation for the inadequacy of *Ddx3y* to compensate for loss of *Ddx3x* lies in gene expression. While *Ddx3x* is ubiquitously expressed, *Ddx3y* expression is much more restricted (30). Indeed, *Ddx3y* transcripts are at least 4-fold less abundant in small pre B cells than at any other stage of B cell development (ImmGen RNA-seq Gene skyline, (21)), including early pro B cells. This illuminates the potential for ample *Ddx3y* in pro B cells to counterbalance loss of *Ddx3x* at this stage, while much lower expression of *Ddx3y* in pre B cells is inadequate. There are reports of compensatory elevations in *Ddx3y* expression in the absence of *Ddx3x (31, 32)*, which could also act to offset loss of *Ddx3x*. Alternatively, while DDX3X and DDX3Y are suggested to possess redundant biochemical activity in translation (17), distinct functional capacities of these helicases to regulate RNA metabolism exist that could prevent *Ddx3y* rescue of *Ddx3y*-deficient B cells (18).

Nevertheless, the B cells present in mice lacking *Ddx3x* are not ‘normal’, suggesting that Ddx3x may be required for specific aspects of B cell differentiation in males. For example, peripheral B cells in hemizygous *Ddx3x*-deficient mice predominately exhibit a marginal zone phenotype. This could reflect *Ddx3x*-dependent changes in mature B-cell intrinsic expression of microRNAs (e.g., miR-146a), transcription factors (e.g., ARID3A/Bright), or receptors (e.g., Notch2) that play defined roles in fate decisions between the marginal zone and follicular B cell lineages (33-36). High levels of B cell-activating factor (BAFF) could also favor marginal zone differentiation. Of note, we observe elevated BAFF in sera of *Ddx3x*^*ΔVav1*^ male mice (control = 9.5 ± 1.1 ng/mL, *Ddx3x*^*ΔVav1*^ =22.2 ± 4.3 ng/mL, n=4, p=0.0075 unpaired t-test with Welch’s correction). Importantly, marginal zone phenotype skewing could also be a consequence of B-cell receptor signal strength and altered selection of B cells during development (37). Consistent with this possibility, we observed significant skewing of the ratio of κ to λ light chain usage in peripheral B cells. These shifts have the potential to impact humoral immunity, as the exaggerated lymphopenia linked to hematopoietic deletion of *Ddx3x* (*Vav1-Cre*) resulted in selective hypergammaglobulinemia. Lymphopenia-driven homeostatic proliferation of T and B cells in these mice was the apparent driver of increased IgG1 and IgG2b expression by marginal zone B cells. However, the elevated levels of IgM, IgG2b, and IgG1 in *Ddx3x*^*ΔVav1*^ mice were not mirrored in *Ddx3x*^*ΔMb1*^ mice, suggesting that hypergammaglobulinemia is driven by B cell extrinsic mechanisms.

Consistently but to varying degrees that depend on the sex of mice and nature of *Cre*-drivers, B cell development is throttled by ablation of *Ddx3x* at stages associated with cell cycle arrest, increased epigenetic accessibility of immunoglobulin genes, and DNA recombination (38). We made numerous attempts to ascertain mechanisms by which *Ddx3x* regulates these events at the early pro B and small pre B stages of B-cell differentiation, many of which were less than ideally revealing. Nevertheless, examination of male pre B cells revealed a role for *Ddx3x* in supporting efficient κ light chain rearrangement that is reflected by skewing light chain usage in peripheral B cells. This was putatively linked to dysregulated expression of the histone reader *Brwd1*, which is required to remodel chromatin at the Igκ locus to make this DNA accessible to recombination machinery (27, 39, 40). Indeed, the perturbations in bone marrow B cell development in *Ddx3x*-deficient male mice largely mirror defects seen in mice lacking *Brwd1* (27). The mechanism by which DDX3X regulates *Brwd1* expression remains ill-defined, but could involve direct interaction between DDX3X protein and *Brwd1* transcripts that facilitate known functions of DDX3X in RNA metabolism (6). DDX3X could also indirectly modulate expression of RNA or protein mediators that in turn regulate BRWD1 expression.

Alternatively, repression of *Brwd1* expression and immunoglobulin light chain recombination could be a consequence of failed cell cycle arrest in *Ddx3x*-deficient pre B cells. Indeed, *Brwd1* is repressed by IL-7 receptor signaling (27), where IL-7 promotes ongoing proliferation of developing B cells prior to cell cycle arrest at the pre B cell stage. *Ddx3x* can influence the expression levels of cyclins, including cyclin E1 (41), which are critical for control of cell cycle progression. Exaggerated proliferation of pre B cells in *Ddx3x*^*ΔVav1*^ mice is associated with elevated expression of *Ccnd2* (cyclin D2), the overexpression of which can promote aberrant cellular proliferation (42). We also observed overexpression of *Ccna2* (cyclin A2), Cdc25a, and Cdk6 in male *Ddx3x*^*ΔVav1*^ pre B cells (**Supplemental Table 1**), consistent with sustained proliferation. Thus, failure to properly exit the cell cycle is another likely contributor to reduced *Brwd1* upregulation and light chain recombination at the pre-B cell stage.

In addition to potentially supporting *Brwd1* expression and cell cycle exit, our study does not rule out possible roles for *Ddx3x* in immunoglobulin gene rearrangement or cell survival. Indeed, there is evidence that DDX3X is actively recruited to sites of DNA damage, interacts with DNA repair proteins, and plays a role in both homologous recombination and non-homologous end-joining repair pathways (43). These types of DNA damage repair are essential for resolving the double strand DNA breaks generated during immunoglobulin gene recombination (38). Unfortunately, we did not enrich the relevant immunoglobulin gene loci undergoing recombination in pro B (*Igh* locus) or pre B (*Igk* locus) in anti-DDX3X chromatin immunoprecipitations from sorted wild-type bone marrow B cell subsets (data not shown). A second possibility is that *Ddx3x* is important for survival of developing pre B cells. Indeed, *Ddx3x* is implicated in pro-survival signaling in other lymphocytes, including NK cells (25), though we have not observed increased apoptosis of *Ddx3x*-deficient B cells in vitro or in vivo (data not shown).

In summary, we use distinct *Cre-*driven editing strategies to strengthen the existing premise that *Ddx3x* plays a role in B cell development. Our data demonstrate a more stringent requirement for *Ddx3x* in female B cells than revealed by prior research. Moreover, we describe DDX3-regulated checkpoints at select stages of B cell development that are related to *Ddx3x*-dependent cell cycle arrest, epigenetic, and transcriptional changes that are required for efficient κ light chain recombination. These mechanisms are consequential for κ/λ light chain repertoire diversity and balanced differentiation of follicular verus marginal zone B cells in the periphery, which subsequently impacts immunoglobulin responses caused by inflammation.

## Materials and Methods

### mice

*Ddx3x* floxed mice (*Ddx3x*^*FL/FL*^) were created by Dr. Josef Penninger (Vienna, Austria) (23). These mice were backcrossed to the C57BL/6 background for eight generations and then bred with *Vav1-Cre* mice (MGI:3765313) that were a gift from H. Leighton Grimes. Experiments made use of male hemizygous deficient mice (*Ddx3x*^*FL/Y*^*Vav1-Cre*^*+*^ *or Ddx3x*^*ΔVav1*^) and wild-type (WT) littermate controls (*Ddx3x*^*FL/Y*^*Vav1-Cre*^*neg*^ or *Ddx3x*^*WT/Y*^*Vav1-Cre*^*+/-*^). In other experiments, *Ddx3x*^*FL/FL*^ mice were bred to *Mb1-Cre* (26) mice obtained from Jackson Labs (stock #020505). Subsequent experiments compared male *Ddx3x*^*FL/Y*^*Mb1-Cre*^*+/-*^ (*Ddx3x*^*ΔMb1*^) mice to controls that included *Ddx3x*^*WT/Y*^*Mb1-Cre*^*+/-*^ or *Ddx3*^*FL/Y*^ *Mb1-Cre*^*neg*^ littermates. Female *Ddx3x*^*FL/FL*^*Mb1-Cre*^*+/-*^ (*Ddx3x*^*ΔMb1*^) mice were compared to *Ddx3x*^*FL/FL*^*Mb1-Cre*^*neg*^ (Control) littermates. Mice were housed in a specific pathogen-free barrier facility, and experiments were conducted with approval from the Cincinnati Children’s Institutional Animal Care and Use Committee. Mice were typically used in experiments at 6 to 12 weeks of age, unless otherwise specified.

### Tissue harvest and cell isolation

To examine bone marrow cells, mouse femora and tibias were flushed with 1× PBS (Corning) using a 10-mL syringe and a 26-gauge 3/8-inch needle (BD Worldwide). We filtered the resulting cell suspension through a 70μm filter mesh (Corning). Mouse spleens and lymph nodes were harvested and mashed on top of the 70μm filter mesh. Red blood cells were lysed using an ACK lysing buffer (Thermo Fisher).

### Cell sorting

Cells were stained with fluorochrome-labeled antibodies in 100μL flow buffer (2% fetal bovine sera in PBS) for 30 minutes at 4°C, then fixed with 100μL fixation buffer for 30 minutes at 4°C. Bone marrow small pre-B cells (CD3^-^ Ly6C^-^ Ter119^-^ CD11b^-^ Gr-1^-^ B220^+^ CD43^-^ IgM^-^ IgD^-^) and splenic IgM^+^ B cells (CD3^-^ CD19^+^ IgM^hi^) were sorted using a FACSAria II (BD Biosciences).

### Flow cytometry

Cells were first treated with Fc block (BD Biosciences, 2.4G2) for 5 minutes and then stained with fluorochrome-labeled antibodies in 100μL flow buffer (2% fetal bovine sera in PBS) for 30 minutes at 4°C. Cell were then fixed with 100μL BD Cytofix buffer for 30 minutes at 4°C or permeabilized with BD Cytofix/Cytoperm buffer for intracellular staining for IgM. Data were acquired on LSR II or Fortessa cytometers and analyzed by FACSDiva (BD Biosciences) and FlowJo (TreeStar). Antibodies from BioLegend include those specific for Ter119 (TER-119), CD11b (M1/70), Gr1 (RB6-8C5), B220 (RA3-6B2), CD3ε (500a2), CD4 (GK1.5), CD5 (53-7.3), CD8α (53-6.7), CD19 (6D5), Ly6C (HK1.4), CD24 (M1/69), IgD (11-26c.2a), CD23 (B3B4), CD93 (AA4.1), IgM (RMM-1), NK1.1 (PK136), Ki67 (11F6), CD11a (2D7), CD44 (IM7), CD62L (MEL-14), CD69 (H1.2F3), IgG2b (RMG2b-1), IgG1 (RMG1-1), κ light chain (RMK-45), λ light chain (RML-42), and CD1d (1D1). Antibodies specific for c-Kit (2B8), CD21 (4E3), and Live/Dead reagents were obtained from eBioscience/ThermoFisher. Antibodies specific for CD3ε (145-2C11), CD43 (S7), and BP-1 (BP-1) were obtained from BD Biosciences.

### Proliferation assessment with BrdU

Mice were given an intraperitoneal injection of 50 g/kg BrdU (Sigma-Aldrich) diluted in phosphate buffered saline. After 2.5 hours, bonne marrow was collected and prepared as described above. After staining of surface antigens, cells were subjected to intracellular staining for BrdU using staining kit from eBioscience according to manufacturer’s protocol.

### ELISA

Blood was collected by cardiac puncture using a 1-mL syringe and 26-gauge 3/8-inch needle (BD Worldwide) and centrifuged at 300 x g for 10 minutes. Supernatant was aliquoted and stored at -80°C. Mouse IgA, total IgG, IgM, IgE, IgG1, IgG2c, IgG2b or IgG3 measured using ELISA Ready-SET-Go kits according to manufacturer’s protocols (eBioscience).

### Autoantibody measurements

Anti-dsDNA antibodies were quantified by ELISA on polyvinyl microtiter plates (Costar) coated with poly-L-lysine (1 µg/ml, Sigma-Aldrich, 1 hour at room temperature) and dsDNA (2.5 µg/ml dsDNA in borate-buffered saline overnight at 4°C). After washing and blocking with 3% BSA in borate-buffered saline containing 0.4% Tween-80 for 2 hours at 4 °C, various dilutions of serum samples were added and incubated overnight at 4 °C. After additional washes, plates were incubated with alkaline phosphate–conjugated AffiniPure goat anti-mouse IgG (Fcγ fragment specific, Jackson ImmunoResearch Laboratories) prior to incubation with p-nitrophenyl phosphate liquid substrate (Sigma-Aldrich) for 30 minutes at 37°C and read using the BioTek Epoch ELISA reader (Molecular Devices).

### Microarray analysis

We isolated total cellular RNA from sorted small pre-B cells using the mirVana™ miRNA Isolation Kit (Thermo Fisher Scientific). RNA concentration and integrity were measured on an Agilent Bioanalyzer. Ovation Pico WTA kit (NuGen) was used to amplify and label RNA library. The RNA libraries were evaluated using GeneChip Mouse Gene 2.0 arrays (Affymetrix). RNA library preparation and GeneChip assays were performed by Cincinnati Children’s Gene Expression Core. Raw data were used for analysis with GeneSpring GX software (v13.0, Agilent Technologies). We first applied standard Robust Multi-array Average normalization and baseline transformation to all samples (n=4/group), then removed noise by applying a filter requiring a raw expression value >50. Unpaired t-tests were performed, and p<0.05 and fold change >1.5 were set as benchmarks for differential gene expression between experimental groups. A hierarchical clustered heatmap of the top 20 up-regulated and top 15 down-regulated genes were made using R software (64bit, v3.2.2). Genome data have been deposited in GEO (https://www.ncbi.nlm.nih.gov/geo/) under accession number GSE112549.

### Genotyping PCR

Genomic DNA was extracted from tail clips using the HotSHOT DNA extraction protocol (Bento Bioworks). PCR was performed using the following primers:

*Ddx3x* forward #1 [CAG TAA ATG TGG GAG AAC AGG AAT TAT TC]

*Ddx3x* forward #2 [TGC CAG AAG AAA GCA GTG GAT CTC]

*Ddx3x* reverse [AAA GCT ATC TAG TTC TGA TTG TCG ATA CAT C]

#### Quantitative PCR analysis

We isolated total cellular RNA from sorted small pre-B cells and splenic IgM^+^ B cells using mirVana Kits. We performed reverse transcription using PrimeScript™ RT reagent Kit (Clontech). For qPCR, 10μL SYBR Green master mix (SYBR Premix Ex Taq™ II, Clontech), 0.4μL Rox dye, 1.6μL of 10μM primer mix, 6μL water, and 2μL cDNA template were used in a total 20μL system per sample. Samples were run in triplicates on ABI 7500 Real-Time PCR System (Applied Biosystems). Target genes were normalized to actin or β2-Microglobulin (B2M) mRNA expression levels. Forward and reverse primers used for quantitative PCR (qPCR) are as follows:

*Ddx3x*-F 5’-ACCCCTATCCCAAACTGCAT-3’

*Ddx3x*-R 5’-TCATGACTGGAATGGCTTGT-3’

*Actin*-F 5’- ATGCTCCCCGGGCTGTAT-3’

*Actin*-R 5’- CATAGGAGTCCTTCTGACCCATTC-3’

*Brwd1*-F 5’-TTGCTTCTGGCAGTGGGATTT-3’

*Brwd1*-R 5’-GCTTTCAAGCTCGGCGATTT-3’

*B2m*-F 5’-AGACTGATACATACGCCTGCA-3’

*B2m*-R 5’-GCAGGTTCAAATGAATCTTCA-3’

*Igκ*-Germline-F 5’-GAGGGGGTTAAGCTTTCGCCTACCCAC-3’

*Igκ*-Germline-R 5’-GTTATGTCGTTCATACTCGTCCTTGGTCAA-3’

*degVκ* 5’-GGCTGCAGSTTCAGTGGCAGTGGRTCWGGRAC-3’

*κ-J1-R* 5’-AGCATGGTCTGAGCACCGAGTAAAGG-3’.

#### PCR analysis of Igκ rearrangement

We performed semi-quantitative PCR assays with reverse transcribed cDNA using an updated protocol (27). Instead of genomic DNA, we performed PCR assays from cDNA using *Vk-FW* (AGCTTCAGTGGCAGTGGRTCWGGRAC) and *Ck* (CTTCCACTTGACATTGATGTC) primers, which target rearranged *Vκ-Jκ* transcripts. We sorted small pre-B cells from *Ddx3x*^*ΔMb1*^, *Ddx3x*^*ΔVav1*^, and corresponding control mice. cDNA from WT splenic IgM^+^ B cells were used as a positive control. *B2m* was amplified as an internal control. PCR products were evaluated on a 1% agarose gel and quantified using ImageLab software (BIO-RAD). The intensity of the band for each rearrangement product (*Vκ-Jκ1/2/4/5*) was divided by that of the corresponding *B2m* band, and the resulting value was then normalized to the values obtained from IgM^+^ B cells.

#### Statistical analysis

Experiments were analyzed using GraphPad Prism (Version 10). Individual data points and means are presented. Normal distribution of data was assessed using the D’Agostino & Pearson test or the Shapiro-Wilk test (when n was low). Normally distributed datasets were compared using an unpaired two-tailed t test with Welch’s correction added when variance was unequal; ordinary one-way ANOVA with multiple comparison tests corrected for false discovery rate using the two-stage linear step-up procedure of Benjamini, Krieger and Yekutieli; or Brown-Forsythe ANOVA test (when variance was not equal) with Dunnett’s T3 multiple comparisons test correction. Non-normally distributed data were analyzed using Mann-Whitney test or Kruskal-Wallis test. One sample t-test was used for comparisons where one dataset was normalized to a value of 1. Statistical significance thresholds are defined in legends for figures that denote comparisons above this threshold as ns, and actual p values are shown or ranges are defined in figure legends.

## Acknowledgements

All flow cytometric data were acquired using equipment maintained by the Research Flow Cytometry Facility in the Division of Rheumatology at Cincinnati Children’s Hospital Medical Center. The authors thank the Cincinnati Children’s Comprehensive Rodent and Radiation Facility (RRID:SCR_022624) for assistance with bone marrow chimera generation, and the Genomics Sequencing Facility (RRID:SCR_022630) for assistance with microarray assays. We thank H. Leighton Grimes for provision of Vav1-Cre mice. We are indebted to Harinder Singh for his critical insights and suggestions. All flow cytometric data were acquired using equipment maintained by the Cincinnati Children’s Research Flow Cytometry Facility (RRID: SCR_022635).

## Author contributions

Conceptualization and design of study [KL (equal), SNW (equal), RHS (supporting), JBH (supporting)], project supervision [SNW (lead), JBH (supporting)], acquisition of funding (SNW (equal), JBH (equal), MTW (supporting), LCK (supporting)], performance of experiments and data acquisition [KL (equal), DK (equal), JT (supporting), EM (supporting), HS (supporting), EMK (supporting), OAD (supporting), WS (supporting)], data analysis [KL (equal), DK (equal), RAK (supporting), DR (supporting), JT (supporting), OAD (supporting), MTW (supporting), LCK (supporting), WS (supporting), MM (supporting), SNW (equal), JBH (supporting)], drafting of the manuscript [KL (equal), DK (equal), SNW (equal), JBH (supporting)], provision of reagents and expertise [TP (supporting), MF (supporting), MM (supporting), DR (supporting), RHS (supporting), JMP (supporting)], and critical editing of the manuscript [JT (supporting), EM (supporting), HS (supporting), EMK (supporting), RAK (supporting), OAD (supporting), TP (supporting), MF (supporting), MM (supporting), MTW (supporting), LCK (supporting), WS (supporting), DR (supporting), RHS (supporting), JMP (supporting)].

